# Utilizing tACS to enhance memory confidence and EEG to predict individual differences in brain stimulation efficacy

**DOI:** 10.1101/2024.05.27.596015

**Authors:** Syanah C. Wynn, Tom R. Marshall, Erika Nyhus

## Abstract

The information transfer necessary for successful memory retrieval is believed to be mediated by theta and gamma oscillations. These oscillations have been linked to memory processes in electrophysiological studies, which were correlational in nature. In the current study, we used transcranial alternating current stimulation (tACS) to externally modulate brain oscillations to examine its direct effects on memory performance. Participants received sham, theta (4 Hz), and gamma (50 Hz) tACS over frontoparietal regions while retrieving information in a source memory paradigm. Linear regression models were used to investigate the direct effects of oscillatory non-invasive brain stimulation (NIBS) on memory accuracy and confidence. Our results indicate that both theta and gamma tACS altered memory confidence. Specifically, theta tACS seemed to lower the threshold for confidence in retrieved information, while gamma tACS appeared to alter the memory confidence bias. Furthermore, the individual differences in tACS effects could be predicted from electroencephalogram (EEG) measures recorded prior to stimulation, suggesting that EEG could be a useful tool for predicting individual variability in the efficacy of NIBS.

## 1. Introduction

Memory is an essential part of cognition, which we do not solely use when studying for a test but is an integral part of our lives. The latter is evidenced by the severe debilitating effects memory loss has on people. Because of this, memory has been a popular topic in the field of cognitive neuroscience, which has provided us with information about the brain mechanisms involved. It is now widely accepted that memories are not retrieved by a single brain region, but by a network of brain regions, including the medial temporal lobe (MTL), dorsolateral prefrontal cortex (DLPFC) and the posterior parietal cortex (PPC) (Bastin et al., 2019; Benoit & Schacter, 2015; Nilakantan, Bridge, Gagnon, VanHaerents, & Voss, 2017; Wang et al., 2014). The integrative memory model (Bastin et al., 2019) proposes that regions in the MTL are part of the ‘core system’, which stores representations of specific memories, forming the content of a memory. The DLPFC is part of the ‘attribution system’, which guides memory search and monitors retrieved information by cross-referencing the current task context with the retrieved information. This processing is then translated into a decision about the retrieved information and a subjective memory experience (“I am very confident this retrieved information is correct and relevant for the current task”). The PPC is part of the ‘connectivity hub’ which enables transfer of information between the core and attributional system. When we are cued to retrieve memories, these brain regions interact closely to enable us to use the retrieved information to make memory-related decisions (e.g., determining if we remember something correctly) and give us a subjective experience (e.g., feeling sure or unsure about the memory, or that the information is at the tip of your tongue).

Theta (3-7 Hz) and gamma (30-100 Hz) oscillations are thought to play an important role in the information transfer in memory related processes. For example, when comparing recognition of an item in isolation (item memory) to recognition that incorporates the retrieval of contextual information (source memory), the latter was associated with greater frontoparietal functional connectivity. This functional connectivity was found in the low gamma range and was modulated by low theta (Burgess & Ali, 2002). These results seem to concur with the notion of memory-related theta-gamma coupling. In this theta-gamma coupling, each gamma cycle represents a specific memory representation, which is superimposed onto different phases of the theta cycle (Griffiths et al., 2019; Heusser, Poeppel, Ezzyat, & Davachi, 2016; Karlsson, Lindenberger, & Sander, 2022; Lisman & Idiart, 1995; Lisman & Jensen, 2013; Roehri, Brechet, Seeber, Pascual-Leone, & Michel, 2022; Ursino, Cesaretti, & Pirazzini, 2023). If theta and gamma oscillations are important for communication during memory, the power in those frequencies should increase when remembering information. As predicted, theta power over frontal and parietal areas increases during successful item (Chrastil et al., 2022; Duzel et al., 2003; Duzel, Neufang, & Heinze, 2005; Wynn, Daselaar, Kessels, & Schutter, 2019; Wynn, Kessels, & Schutter, 2020b) and source memory (Addante, Watrous, Yonelinas, Ekstrom, & Ranganath, 2011; Gruber, Tsivilis, Giabbiconi, & Muller, 2008; Guderian & Duzel, 2005; Herweg et al., 2016; Wynn, Townsend, & Nyhus, 2024). Literature is limited on neocortical gamma, but there is evidence for an increase in gamma power during both item (Gruber et al., 2008) and source memory (Burgess & Ali, 2002) from previous EEG studies.

Because of this link between oscillatory brain activity and memory, non-invasive brain stimulation (NIBS) methods targeting oscillations, like transcranial alternating current stimulation (tACS) and oscillatory direct current stimulation (otDCS), have been used to modulate episodic memory. In tACS, weak alternating currents in a certain frequency are applied to the head to stimulate the brain. In otDCS, an alternating currents is superimposed onto a direct current, making the current oscillate around a non-zero value (Herrmann, Rach, Neuling, & Struber, 2013). Theta tACS has been utilized to facilitate memory encoding in several studies and they report promising findings (Alekseichuk, Turi, Veit, & Paulus, 2020; Antonenko, Faxel, Grittner, Lavidor, & Floel, 2016; Klink, Peter, Wyss, & Kloppel, 2020; Lang, Gan, Alrazi, & Monchi, 2019). In addition, the effect of offline anodal otDCS at theta frequency on retrieval has been investigated. When targeting the left PPC, theta otDCS has been shown to improve associative memory as compared to sham (Vulic et al., 2021). However, as similar effects were observed when using non-oscillatory tDCS, these effects can not directly be attributed to theta oscillations. Anodal theta otDCS targeting the left DLPFC had no effect on item memory, while it did impair source memory (Mizrak et al., 2018). Both otDCS findings are surprising given the EEG literature on theta oscillations and memory previously discussed. Since both studies did not stimulate during retrieval (‘online’), but right before (‘offline’), this could explain the inconsistencies with the EEG literature. A study that did stimulate during retrieval, found immediate and prolonged effects of left PFC gamma tACS on item memory (Nomura, Asao, & Kumasaka, 2019). However, in this study stimulation was also applied during encoding, making it impossible to disentangle encoding from retrieval tACS effects. Another study showed that medial PFC theta tACS was to be able to improve item memory in people with subjective memory complaints and that this can be a viable intervention for this population (Varastegan et al., 2023). In general, the subjective aspect of memory may be more susceptible to NIBS interventions as it can be seen as a more sensitive measure than objective accuracy from ‘old/new’ judgements. Evidence for this comes from a study that applied theta tACS during retrieval, targeting the PPC bilaterally (Wynn, Kessels, & Schutter, 2020a). In this study item and source memory were not affected by the tACS condition, but the subjective memory experience was reduced.

In the current study, we compared the efficacy of frontoparietal theta and gamma tACS on memory accuracy and confidence, and explored the EEG components that can predict individual differences in this efficacy. Our participants performed a source memory task and received stimulation during retrieval. The target location of stimulation was chosen to match the core regions of the retrieval network. We aimed to improve source memory accuracy and memory confidence by enhancing communication between the DLPFC and PPC, and indirectly the MTL. Based on EEG and fMRI literature, we hypothesized that both gamma and theta tACS would improve item memory, source memory, and memory confidence. Yet, the studies using NIBS to alter purely retrieval-related processes suggest that effects may only be expected on the subjective measure of memory confidence. Given the limited research on the specific role of neocortical gamma and memory, gamma tACS effects were anticipated to be less pronounced, as compared to theta tACS effects. Furthermore, the significant tACS effects were explored further by using EEG to predict individual differences in the efficacy of the brain stimulation.

## 2. Methods

### 2.1. Participants

Fifty-four healthy adult right-handed volunteers (32 females, 22 males) with a mean age of 20 (*SD* = 1.55) were included in this study. Four participants were replaced to maintain our intended sample-size of 54, due to not adhering to task instructions (N=2) and failure to complete all experimental sessions (N=2). All had normal or corrected-to-normal vision, were fluent English speakers, right-handed and free from self-reported neurological or psychiatric conditions. Main exclusion criteria were skin disease, metal in their cranium; epilepsy or a family history of epilepsy; history of other neurological conditions or psychiatric disease; heart disease; use of psychoactive medication or substances; pregnancy. Stimulation parameters were in agreement with the International Federation of Clinical Neurophysiology safety (Rossi, Hallett, Rossini, Pascual-Leone, & Safety of TMS Consensus Group, 2009). The study was approved by the institutional review board of Bowdoin College, Brunswick, USA, and carried out in accordance with the standards set by the Declaration of Helsinki.

### 2.2. Stimuli

Stimuli were presented on a personal computer screen with a 21-inch monitor. Stimulus presentation and recording of responses were attained using E-Prime 2.0 software (Psychology Software Tools, Pittsburg, PA). The stimulus material consisted of 400 words per session, varying per participant, randomly chosen from a pool of 1778 words, selected from the MRC Psycholinguistic Database (http://websites.psychology.uwa.edu.au/school/MRCDatabase/uwa_mrc.htm). All words in this database are scored on word frequency, familiarity, and concreteness, which combined leads to an ‘imageability’ rating between 100 and 700 (Coltheart, 1981). We only included nouns and adjectives that had an imageability rating of >300. For each session, parallel versions of word lists that were equated on imageability (*M* = 507), familiarity (*M* = 509), word frequency (*M* = 54), number of letters (*M* = 6) and word type (91-93% nouns) were used in the experiment. For each session, the words used for encoding and retrieval were randomly generated separately for each participant.

### 2.3. TACS parameters

TACS was delivered by a battery-driven constant DC current stimulator (NeuroConn, DC-Stimulator Plus, neuroConn GmbH, Ilmenau, Germany) using three electrodes (2 × 9 cm^2^, 1 × 35 cm^2^) at a 4 or 50 Hz alternating current intensity of 2 mA (peak-to-peak) for maximally 30 minutes. There was a ramp up and ramp down period of 10 seconds, in which the intensity was gradually increased or decreased between 0 and 2 mA (peak-to-peak). During sham tACS, the 10-second ramp up period was followed by 30 seconds of real stimulation, after which the intensity was ramped down in 10 seconds to 0 mA. Impedance was kept under 15 kΩ throughout the experiment, with a mean impedance at the start of stimulation of 9.43 kΩ (*SD* = 1.00). TACS was administered via two active electrodes over AF4 and P5 electrode sites conforming to the International 10-20 system, targeting the right DLPFC and the left PPC (size: 9 cm^2^, current density: 0.11 mA/cm^2^). These regions were chosen based on previous EEG and fMRI literature implicating them as major regions of interest (Nyhus & Badre, 2015; Nyhus, Engel, Pitfield, & Vakkur, 2019; Vilberg & Rugg, 2008). The reference electrode was centred over Cz (size: 35 cm^2^, current density: 0.06 mA/cm^2^). Conductive adhesive paste was used to enhance conductivity between the electrodes and the scalp and to hold the electrodes in place.

To estimate the electric field density and distribution of this setup, a simulation was performed on a standard brain using SimNIBS (Opitz et al., 2015) (see Figure 1). The specific theta frequency (4 Hz) was based upon a previous study using tACS to alter memory (Wynn et al., 2020a). The specific gamma frequency (50 Hz) was selected based upon: (1) non-overlapping harmonics with the target frequency, (2) a tACS sinus that has a period that fits in an integer number of EEG samples, (3) a frequency that does not overlap with the rhythm of heartbeat (∼1 Hz) and respiration (∼0.16 - 0.33 Hz), and (4) a frequency that does not overlap with frequencies that can induce phosphenes (∼7 - 30 Hz) (Kanai, Chaieb, Antal, Walsh, & Paulus, 2008).

**Figure 1.**
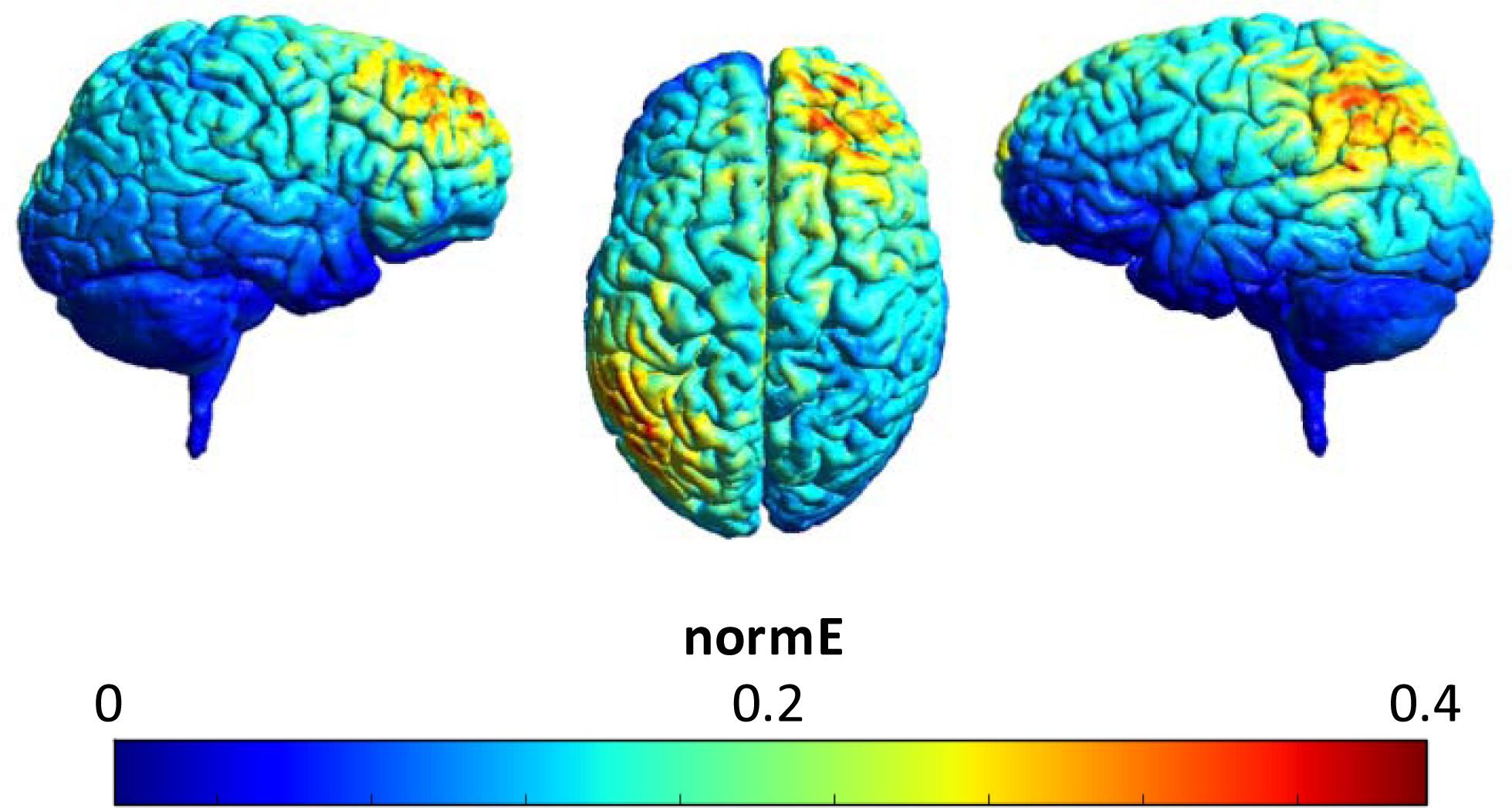
Simulated electric field distribution of stimulation targeting the R-DLPFC and L-IPC, with the use of the SimNIBS software (Opitz, Paulus, Will, Antunes, & Thielscher, 2015).

### 2.4. Procedure

All participants received written and oral information prior to participation but remained naive regarding the aim of the study. Each volunteer provided written informed consent at the beginning of the first session. In this first session, participants did not receive any stimulation, and only EEG was recorded. In the following sessions, participants received theta (4 Hz) tACS, gamma (50 Hz) tACS, or sham (4 Hz) tACS across three sessions in a counterbalanced order. The four sessions were scheduled to be separated by exactly one week and controlled for time of day. Electrodes were placed at the start of each experimental session.

In the intentional encoding phase of the memory task, trials began with the presentation of solely the response options on the bottom of the screen for 80-120 ms (jittered; see Figure 2). Throughout the trial, these response options remained on the screen. Next, the task cue (‘Place’ or ‘Pleasant’) was presented in the middle of the screen in yellow font for 500 ms, followed by a blank mask for 200 ms and the presentation of the to-be-encoded capitalized word for 500 ms. The cue informed the participants on the encoding task in the current trial. When the cue ‘Place’ was presented, participants had to conjure up an image of a scene of a spatial environment that relates to the word that was presented right after. For example, for the word ‘dirty’, they could imagine a dirty scene or place, such as imagining a scene of a garbage dump or a messy room. When the cue ‘Pleasant’ was presented, participants had to pay attention to the meaning of the word that was presented right after and evaluate the pleasantness of the word. For example, for the word ‘dirty’, they could imagine that it is ‘unpleasant’. Following the word offset, participants had 4 seconds to perform the encoding task while a fixation cross was presented on screen. Thereafter, a question mark replaced the fixation cross, and they had 700 ms to indicate how successful they were at completing the encoding task by responding on a keyboard with their dominant, right hand: ‘H’ = unsuccessful, ‘U’ = partially successful, ‘I’ = successful. In total there were 200 trials; 100 words encoded during the place task and 100 words encoded during the pleasantness task.

**Figure 2.**
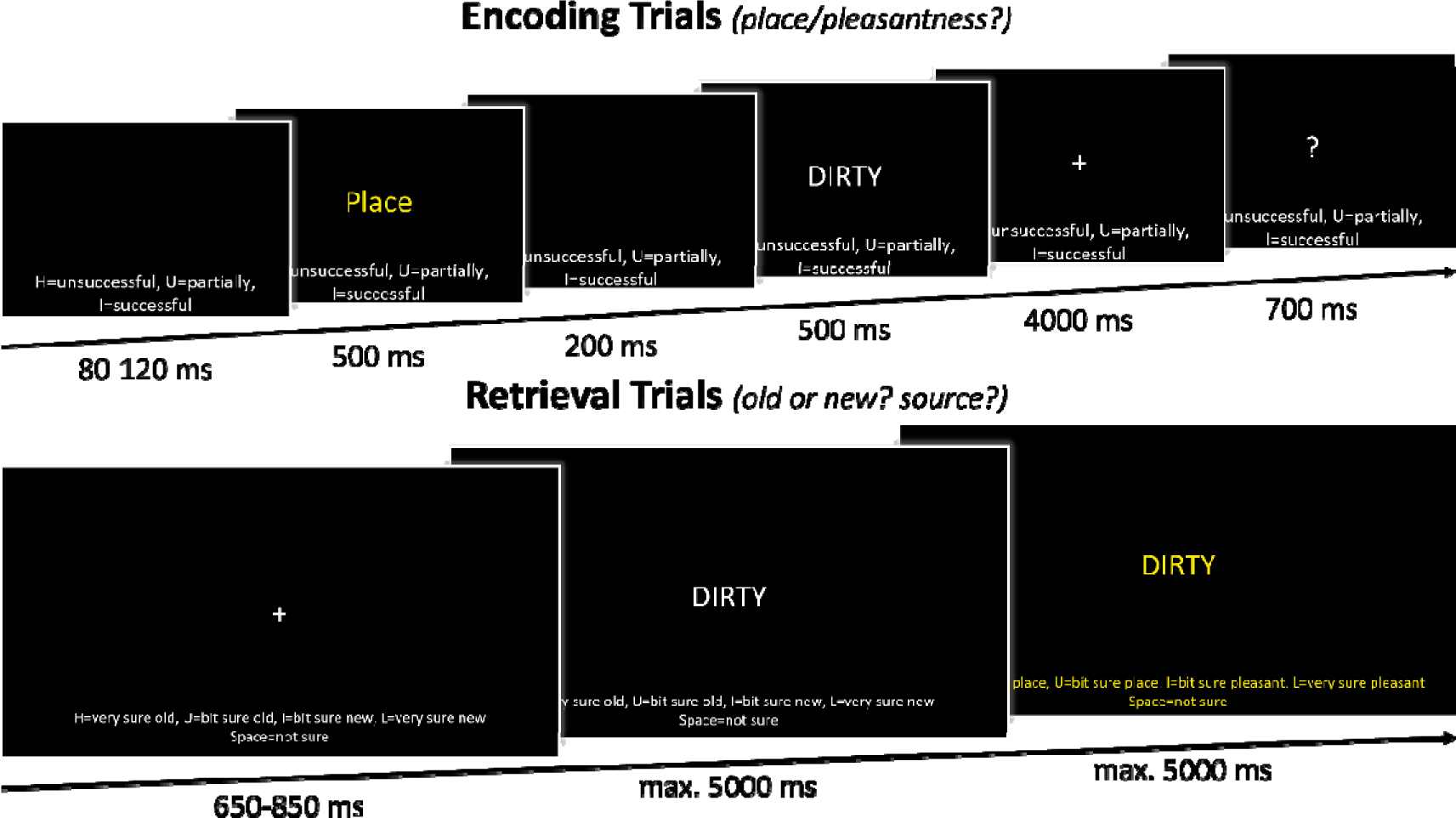
Schematic overview of the memory task. In the encoding phase, participants either had to imagine a spatial scene (place task) or rate the pleasantness (pleasantness task) regarding the presented word. They indicated how successful they were in completing this encoding task. In the retrieval phase, participants first made an ‘old/new’ response. In the case of an ‘old’ response, participants were asked to indicate which encoding task was performed when first encountering that word.

After the encoding task, participants completed a math task to diminish any rehearsal and recency effects. In this task, mathematical equations (e.g., 56 + (5 + 5) × 2 – 30) were presented on the computer screen for 15 minutes. Participants were informed that this task was a distraction task, that they should not be nervous about it and just try their best. The math task was self-paced, and participants could alter their answer prior to responding. Responses were attained using the numbers on the keyboard.

TACS was delivered during retrieval, with the onset one minute before the participants started the experimental retrieval trials. Participants received tACS in one of the three conditions while performing the recognition task, including the 200 ‘old’ words that were presented during encoding and 200 ‘new’ words (see Figure 2). Like the encoding trials, the response options were presented on the bottom of the screen throughout the trial. The retrieval trials started with a 650-850 ms (jittered) presentation of a fixation cross. This was followed by a centrally presented capitalized word. Participants were instructed to indicate whether they thought the word was ‘old’ or ‘new’, considering the confidence they had in their decision, on a 5-point scale. Responses were given on a keyboard with their dominant, right hand: ‘H’ = very sure old, ‘U’ = bit sure old, ‘space bar’ = not sure, ‘I’ = bit sure new, ‘L’ = very sure new. Participants had 5 seconds to submit their response and their response immediately advanced the trial. However, to ensure a sufficient time frame for EEG analyses, within the first 800 ms of word presentation, entering a response would not advance the trial immediately. When their response was ‘not sure’, ‘bit sure new’, or ‘very sure new’ the next trial started after their ‘old/new’ decision was finalized. When their response was ‘bit sure old’ or ‘very sure old’, a new screen was presented, and participants could indicate the remembered encoding source of the word. On this screen the source response options were presented: ‘H’ = very sure pleasant, ‘U’ = bit sure pleasant, ‘space bar’ = not sure, ‘I’ = bit sure place, ‘L’ = very sure place. To make it more salient to the participants that a source decision was now required, both the word in the center of the screen and the response options on the bottom were presented in a yellow font. Participants had 5 seconds to indicate their response on the keyboard, after which the next trial began.

To familiarize participants with the memory task, fifteen practice trials preceded both the encoding and retrieval phase of the experiment. Stimuli used during the practice trials were not used in the experimental trials. Following every 100 experimental trials there was a short break of a minimum of 30 seconds, after which participants had 30 seconds to indicate that they were ready to continue. At the end of the fourth session, volunteers were debriefed and received compensation for participation.

### 2.5. EEG recording and analyses

EEG was recorded throughout the experimental sessions. EEG signals were recorded and amplified with an actiCHamp system (Brain Products, Munich, Germany). During the first session, without tACS, data was recorded from 64 channels, during the following three sessions, data was recorded from 32 channels due to the limited space on the head from the concurrent tACS electrode placement. The amplified analogue voltages (0.1–100 Hz bandpass) were digitized at 10 kHz. As we wanted to investigate how EEG data can be used to predict subsequent tACS effects, here we only analysed and reported on the EEG data from the first session. In addition, it is not possible to look at possible online effects of the tACS on EEG, due to the tACS-induced EEG artifact that is magnitudes larger and in the same frequency window as the brain signal of interest (Noury & Siegel, 2018).

EEG pre-processing and analyses were performed with the use of MATLAB (v2022b, MathWorks Inc., Natick MA) in combination with the FieldTrip toolbox (Oostenveld, Fries, Maris, & Schoffelen, 2011) and the EEGLAB toolbox (Delorme & Makeig, 2004). Raw signals were downsampled to 1000 Hz and re-referenced to an average reference. The data was high-pass filtered at 0.1 Hz, low-pass filtered at 58 Hz, and an additional band-pass filter was used to remove residual line noise at 60 Hz. Subsequently, the retrieval data was epoched into stimulus-locked time windows. The minimum stimulus presentation duration during retrieval was 800 ms. However, since retrieval was self-paced, stimulus presentation duration varied. For this reason, epochs started 500 ms before stimulus onset and ended either 505 ms before the onset of the following stimulus or after 2000 ms. This resulted in the shortest retrieval epoch being -500 to 1045 ms, and the average epoch being -500 to 1810 ms. Epochs with transient muscle or electrode artifacts were rejected based on visual inspection. Additional artifacts were removed using independent component analysis (ICA) in combination with EEGLAB’s ICLabel (Pion-Tonachini, Kreutz-Delgado, & Makeig, 2019). Components classified as muscle artifacts (probability > .9), eye artifacts (probability > .8), heart artifacts (probability > .8), and channel noise artifacts (probability > .9) were removed from the data. A final artifact check was done after ICA by manual inspection.

Spectral power was extracted using a Fourier analysis with a Hanning taper. Data was zero-padded to a total length of 5 seconds and frequencies that were assessed ranged from 1 to 57 Hz in 1 Hz steps. To investigate whether stimulation efficacy was dependent on the match between the peak frequency and the tACS frequency, the peak frequency deviation was calculated. This was done by extracting the frequency with the highest power in a range surrounding the theta and gamma tACS frequencies, 1-7 Hz and 45-55 Hz respectively. This ‘peak deviation’ was defined as the absolute difference between the tACS stimulation frequency (4 or 50 Hz) and the EEG peak frequency in the corresponding frequency band (1-7 Hz or 44-55 Hz). To facilitate the comparison of spectral power and peak frequency deviations between the theta and gamma frequency bands and to minimize the effect of the 1/f activity, the EEG power was baseline-corrected to the -500 to -250 ms time window, with a relative baseline.

As our tACS set-up was designed to increase synchronization in frontoparietal regions, it was of interest to investigate whether an EEG-based measure of rhythmic neuronal synchronization could predict individual differences in stimulation effects. To quantify this synchronization, we used the pairwise phase consistency (Vinck, van Wingerden, Womelsdorf, Fries, & Pennartz, 2010) between the frontal (AF4, AFz, AF8, F2, F4) and parietal (P5, P3, P7, CP5, PO7) channels. In addition, although stimulation was only applied at one frequency at a time, given that coupling between theta and gamma is thought to be important for binding memory elements together (Griffiths et al., 2019; Lisman & Jensen, 2013), we explored whether theta-gamma coupling could predict individual differences in performance enhancement by tACS. This phase-amplitude coupling between theta phase and gamma amplitude was calculated for the frontal and parietal channels.

To prepare the data for further statistical analysis, for each condition, data was averaged separately per frequency band (theta: 1-7 Hz; gamma: 45-55 Hz) and/or channel group (frontal: AF4, AFz, AF8, F2, F4; parietal P5, P3, P7, CP5, PO7). These values were based on the tACS frequencies (4 and 50 Hz), and electrode placement (AF4 and P5) used in the current study.

### 2.6. Statistical analyses

All regression analyses were performed in RStudio (RStudio version 2023.06.2, R version 4.2.0; R Core Team, 2021). To avoid issues with multicollinearity, for each model variance inflation factors (VIFs) were determined. Values greater than 5 indicate that variables are highly correlated and deemed a cause for concern.

For all trial-based behavioral statistical analyses, each trial was coded based on memory status (‘old’ or ‘new’), item/source memory accuracy (‘correct’ or ‘incorrect’), and item/source memory confidence (‘high’ or ‘low’). Due to the low number of ‘a bit sure’ and ‘not sure’ responses, these two responses were combined into one ‘low confidence’ level. For increased readability, where relevant the following memory categories were used: hits (‘old’ and ‘correct’), misses (‘old’ and ‘incorrect’), correct rejections (‘new’ and ‘correct’), and false alarms (‘new’ and ‘incorrect’). For the behavioral analyses, generalized linear mixed effects models were used to test for a significant tACS effect on item/source memory accuracy and confidence, while controlling for confounding behavioral variables (lme4 package (v. 1.1.29); Bates, Mächler, Bolker, & Walker, 2015). A mixed effect model was deemed most appropriate as it can account for within- and between-subject variability, through employing by-participant varying intercepts. Specifically, the following models were used in the analyses:

1. *glmer(Item Accurcy ∼ Stimulation * Memory Status * Con[idence + (1 | Participant)*
2. *glmer(Item Con[idence ∼ Stimulation * Memory Status * Accuracy + (1 | Participant)*
3. *glmer(Source Accurcy ∼ Stimulation * Con[idence + (1 | Participant)*
4. *glmer(Source Con[idence ∼ Stimulation * Accuracy + (1 | Participant)*

As outcome variables were binary, the specific model used was a binomial generalized linear mixed effects model (GLMM). The fixed effects were Stimulation (‘sham’, ‘gamma’, ‘theta’; treatment coding, reference level: ‘sham’), Memory Status (‘old’, ‘new’; sum coding: -1, 1), Accuracy (‘correct’, ‘incorrect’, sum coding: -1, 1), and Confidence (‘high’, ‘low’, sum coding: -1, 1). All binary predictors were sum coded, so that a value of 0 means the middle of the two possible categories. The estimates of the binary predictors thus reflect the main effect of the predictor and half of the difference between the two categories. In other words, the difference between a value of 0, the middle of the categories (the average of the average of the two categories), and a value of 1 (the category that is coded as 1). The predictor Stimulation was treatment coded, so that both active stimulation conditions (‘theta’ and ‘gamma’) were compared to the condition coded as 0, the reference (‘sham’). As our main interest was on the stimulation effects and their possible interaction with memory status, accuracy and/or confidence, we included these interactions in the model. This was additionally justified by comparing the fit of the more complex models above to simpler models with the highest interaction dropped, through an analysis of variance (ANOVA). This ANOVA showed the more complex models were significantly better at capturing the data than the simpler models, so the above specified models were chosen. All models had a by-participant varying intercept to consider individual differences. By-participant varying slopes were not included in these models as we did not expect large interindividual variability on specific fixed effects. In addition, fitting multiple random slopes in one model led to singular fit. Significance of the model outputs were generated by the lmerTest package (v. 3.1.3; Kuznetsova, Brockhoff, & Christensen, 2017), which applies the Satterthwaite method for estimating degrees of freedom, with an alpha level of .05. In the case of a significant interaction involving Stimulation, pairwise comparisons on the estimated marginal means were used to further explore this. As the binomial GLMM is a logistic regression model, the estimates are log odds. To facilitate interpretation of the estimates, these values were transformed to probabilities.

For all EEG statistical analyses, each trial was first coded based on memory status (‘old’ or ‘new’), item/source memory accuracy (‘correct’ or ‘incorrect’), and item/source memory confidence (‘high’ or ‘low’). Subsequently, trials were averaged per unique combination, or condition, of the above (e.g., high-confident, correct, old items). As the data used as predictors in the model was taken from session 1 and the outcome data from session 2-4, trial-based analyses could not be utilized here. For these analyses linear models were used to test which of the EEG components had a unique predictive value on the subsequent stimulation effects (lme4 package (v. 1.1.29); Bates et al., 2015). The outcome measures included in the models pertained to the significant tACS effects from the aforementioned behavioral analysis. Therefore, the EEG statistical analyses were post-hoc or second-level analyses, aimed to shed more light on the elements that can predict individual differences in stimulation effects. The behavioral outcome measure was a single value per participant which reflected the stimulation effect being investigated. This was quantified as the differences score in conditions (e.g., high-confident hits during theta stimulation – high-confident hits during sham stimulation). These behavioral outcomes were normalized to proportions (i.e., high-confident hits divided by all hits, high-confident correct rejections divided by all correct rejections). The EEG components were extracted from the encoding and retrieval data from the first session of the participants. Specifically, the following model formats were used in the analyses:

(5) *lm(Behavioral Stimulation E[[ect ∼ Encoding Power + Encoding Peak Deviation + Encoding Phase Synchronization + Encoding Phase Aplitude Coupling + Retrieval Power + Retrieval Peak Deviation + Retrieval Phase Synchronization + Retrieval Phase Aplitude Coupling)*
(6) *lm(Behavioral Stimulation E[[ect ∼ Retrieval Power + Retrieval Peak Deviation + Retrieval Phase Synchronization + Retrieval Phase Aplitude Coupling)*

If the behavioral stimulation effect pertained to ‘old’ items, encoding data was also included (model 5), and when it only pertained ‘new’ items, only retrieval data was included in the model (model 6). Given the number of predictive variables and no a priori hypotheses about specific interactions, only simple effects were included in the model. Linear predictors were standardized, except for ‘peak deviation’, for facilitation of interpretation of the effects. The distribution of residuals was checked by visual inspection. Outliers were detected by utilizing the Cook’s distance, with a cut-off value of 4 / (number of observations – number of explanatory variables - 1). Significance of the model outputs were generated by the lmerTest package (v. 3.1.3; Kuznetsova et al., 2017), with an alpha level of .05.

## 3. Results

### 3.1. Descriptives of behavioural performance

For the full description of the retrieval behavioral measures, see Table 1. During encoding, participants thought of either a place or the pleasantness regarding the presented word with an average imagining success of 89.3% (*SD* = 8.3). After the fixed four second time window, their average imagining reaction time (RT) was 354 ms (*SD* = 44).

**Tabel 1.** Mean values of behavioral performance during memory retrieval, with the standard deviation in brackets.

|  | No tACS<br>*Mean (SD)* | Sham tACS<br>*Mean (SD)* | Gamma tACS<br>*Mean (SD)* | Theta tACS<br>*Mean (SD)* |
| --- | --- | --- | --- | --- |
| Encoding Success (%) | .874 (.084) | .901 (.080) | .900 (.078) | .898 (.088) |
| Encoding RT (ms) | 377.549 (46.759) | 347.229 (40.561) | 347.923 (45.286) | 344.631 (45.158) |
| Item d-prime | 2.308 (0.594) | 2.184 (0.818) | 2.273 (0.860) | 2.266 (0.911) |
| Item HC Responses (%) | .608 (.213) | .554 (.272) | .598 (.280) | .557 (.268) |
| Item RT (ms) | 1642.103 (308.526) | 1429.641 (299.730) | 1398.242 (289.948) | 1427.596 (284.002) |
| Source d-prime | 1.640 (0.729) | 1.929 (0.782) | 2.009 (1.020) | 2.014 (0.935) |
| Source HC Responses (%) | .544 (.220) | .549 (.232) | .556 (.245) | .546 (.247) |
| Source RT (ms) | 1273.080 (446.980) | 892.937 (360.883) | 875.685 (337.315) | 871.821 (330.500) |

SD = standard deviation, HC = high-confident, RT = reaction time, ms = milliseconds

On average, participants went through 58 (SD = 26.15) math equations with an average response time of 17 (SD = 7.95) seconds and an accuracy of 78 (SD = 15.05) percent. Given their performance, we deem it unlikely that participants were actively rehearsing items during the 20 minutes break between encoding and retrieval.

The average item memory performance, as quantified by d-prime, was 2.26 (*SD* = 0.77) and the average RT, for the old/new judgement was 1474 (*SD* = 296) ms. The average source memory performance, as quantified by d-prime, was 1.90 (*SD* = 0.87) and the average RT for the source judgement was 978 (*SD* = 369) ms.

### 3.2. TACS effects on behavioural memory measures

To look at the effects of tACS on behaviour, models were used to elucidate the effects of tACS session on the behavioural outcome measure. The model estimates reflect probabilities of the binary outcome variable (e.g., Item Memory Accuracy: 0 = incorrect, 1 = correct) having a value of 1 (e.g., the probability of a correct response). Therefore, tACS effects were defined as a difference in ‘predicted probability’ of the outcome measure between the active tACS conditions (‘theta’ and ‘gamma’) and the control/reference tACS condition (‘sham’). To account for tACS effects interacting with other behavioural variables, we looked at interactions involving the variable Stimulation in addition to the main effects of Stimulation.

For item memory accuracy, the predicted probability of making a correct memory decision was influenced by 3-way interactions between ‘Theta tACS’, ‘Memory Status’, and ‘Confidence’, and between ‘Gamma tACS’, ‘Memory Status’, and ‘Confidence’ (see Figure 3 and Table 1). Post-hoc tests (see Table 2) on the estimated marginal means showed that the predicted probabilities of a correct decision for low-confident new, high-confident new, and low-confident old items did not differ significantly between stimulation conditions. However, for high-confident old items, the predicted probability of a correct decision during Theta stimulation was significantly higher than during Sham stimulation and Gamma stimulation. Specifically, based on this model, we expect that in sessions where participants are receiving theta tACS, high-confident hits occur 2.9% and 1.8% more often, as compared to the gamma and sham tACS sessions. Low-confident hits, low-confident correct rejections, and high-confident correct rejections are statistically equally likely to occur during all three stimulation conditions.

**Figure 3.**
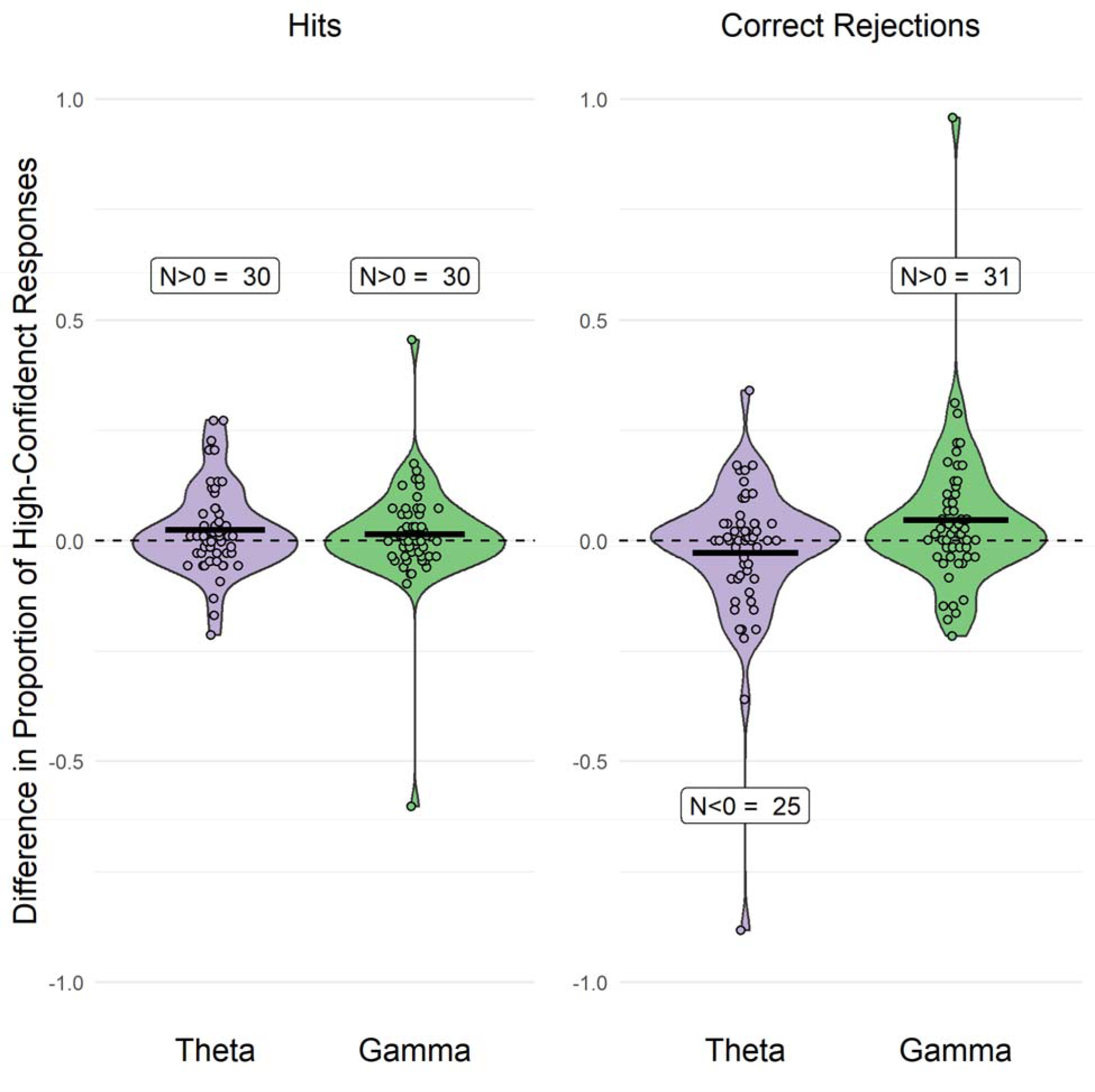
Difference in the proportion of high-confidence hits and correct rejections between sham tACS and the two active stimulation conditions (theta and gamma). The dots represent individual participants, with the number of participants (N) displaying values above or below 0 indicated in the graph. This illustrates the number of participants exhibiting the group-level tACS effect.

**Tabel 1.**
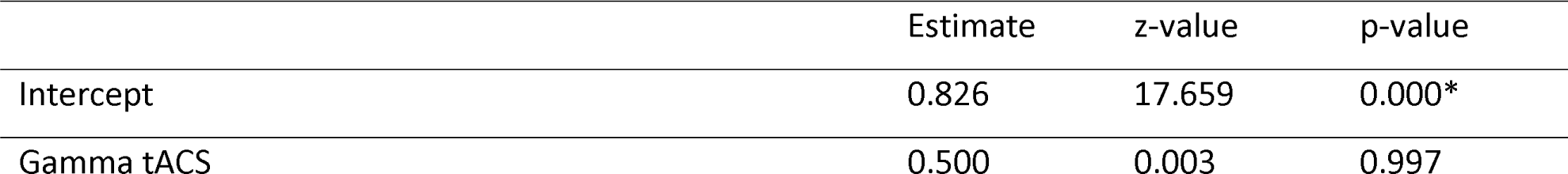
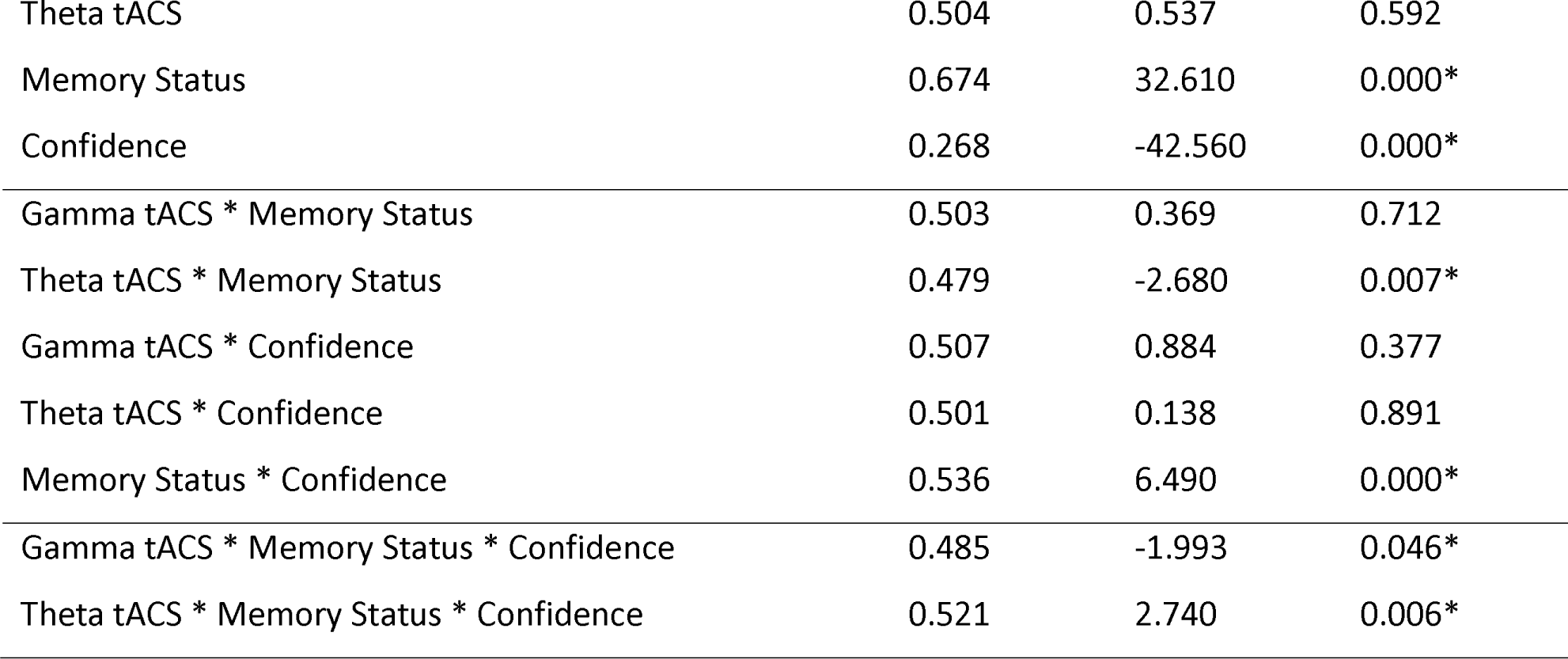
Item Memory Accuracy - tACS Model.

**Table 2.** Item Memory Accuracy - Post-hoc Tests.

| Condition | EM Means 1 |  | EM Means 2 |  | z-ratio | p-value | Eff. size |
| --- | --- | --- | --- | --- | --- | --- | --- |
| Old High Confidence | Theta | 0.897 | Sham | 0.879 | 3.516 | 0.001* | 0.181 |
| Old High Confidence | Gamma | 0.868 | Sham | 0.879 | -2.058 | 0.099 | -0.101 |
| Old High Confidence | Theta | 0.897 | Gamma | 0.868 | 5.604 | 0.000* | 0.283 |
| Old Low Confidence | Theta | 0.425 | Sham | 0.420 | 0.380 | 0.923 | 0.019 |
| Old Low Confidence | Gamma | 0.440 | Sham | 0.420 | 1.560 | 0.263 | 0.079 |
| Old Low Confidence | Theta | 0.425 | Gamma | 0.440 | -1.156 | 0.480 | -0.059 |
| New High Confidence | Theta | 0.952 | Sham | 0.959 | -1.730 | 0.194 | -0.157 |
| New High Confidence | Gamma | 0.961 | Sham | 0.959 | 0.506 | 0.868 | 0.046 |
| New High Confidence | Theta | 0.952 | Gamma | 0.961 | -2.277 | 0.059<sup>+</sup> | -0.203 |
| New Low Confidence | Theta | 0.809 | Sham | 0.806 | 0.489 | 0.876 | 0.023 |
| New Low Confidence | Gamma | 0.802 | Sham | 0.806 | -0.472 | 0.885 | -0.023 |
| New Low Confidence | Theta | 0.809 | Gamma | 0.802 | 0.948 | 0.610 | 0.046 |

\* EM = estimated marginal, Eff. size = Cohen's d. The z-ratio and p-values refer to the difference between EM Means 1 and EM Means 2.

For item memory confidence, the predicted probability of making a high-confident response was influenced by a 3-way interaction between ‘Theta tACS, ‘Memory Status’, and ‘Accuracy’ and a 2-way interaction between ‘Gamma tACS’ and ‘Accuracy’ (see Table 3). Post-hoc pairwise comparisons (see Table 4) showed that the predicted probability of a high-confident response for hits during Theta stimulation was significantly higher than during Sham. Whereas the predicted probability a high-confident response for correct rejections during Theta stimulation was significantly lower than during Sham. Gamma stimulation, as compared to Sham, increased the predicted probability of a high-confident response in all conditions. Specifically, based on this model, we expect that in sessions where participants are receiving theta tACS, high-confident hits occur 2.4% more often, as compared to sham tACS sessions. In addition, in theta tACS sessions, we expect them to make 3.8% and 11.3% less high-confident correct rejections, as compared to sham and gamma tACS, respectively. Whereas in gamma tACS sessions, participants are predicted to have 6.2% more high-confident responses, irrespective of memory status or accuracy, as compared to sham tACS sessions.

**Table 3.** Item Memory Confidence – tACS Model.

|  | Estimate | z-value | p-value |
| --- | --- | --- | --- |
| Intercept | 0.469 | -0.540 | 0.589 |
| Gamma tACS | 0.577 | 8.424 | 0.000* |
| Theta tACS | 0.511 | 1.238 | 0.216 |
| Memory Status | 0.337 | -25.664 | 0.000* |
| Accuracy | 0.266 | -38.315 | 0.000* |
| Gamma tACS * Memory Status | 0.510 | 1.138 | 0.255 |
| Theta tACS * Memory Status | 0.496 | -0.450 | 0.653 |
| Gamma tACS * Accuracy | 0.519 | 2.107 | 0.035* |
| Theta tACS * Accuracy | 0.502 | 0.197 | 0.844 |
| Memory Status * Accuracy | 0.567 | 10.270 | 0.000* |
| Gamma tACS * Memory Status * Accuracy | 0.491 | -0.979 | 0.328 |
| Theta tACS * Memory Status * Accuracy | 0.541 | 4.490 | 0.000* |

**Table 4.**
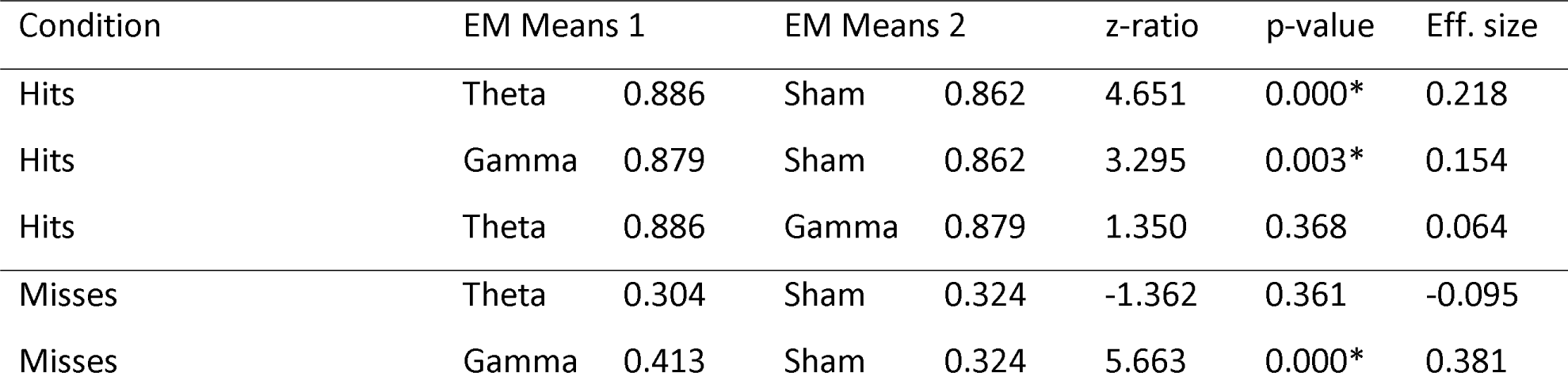
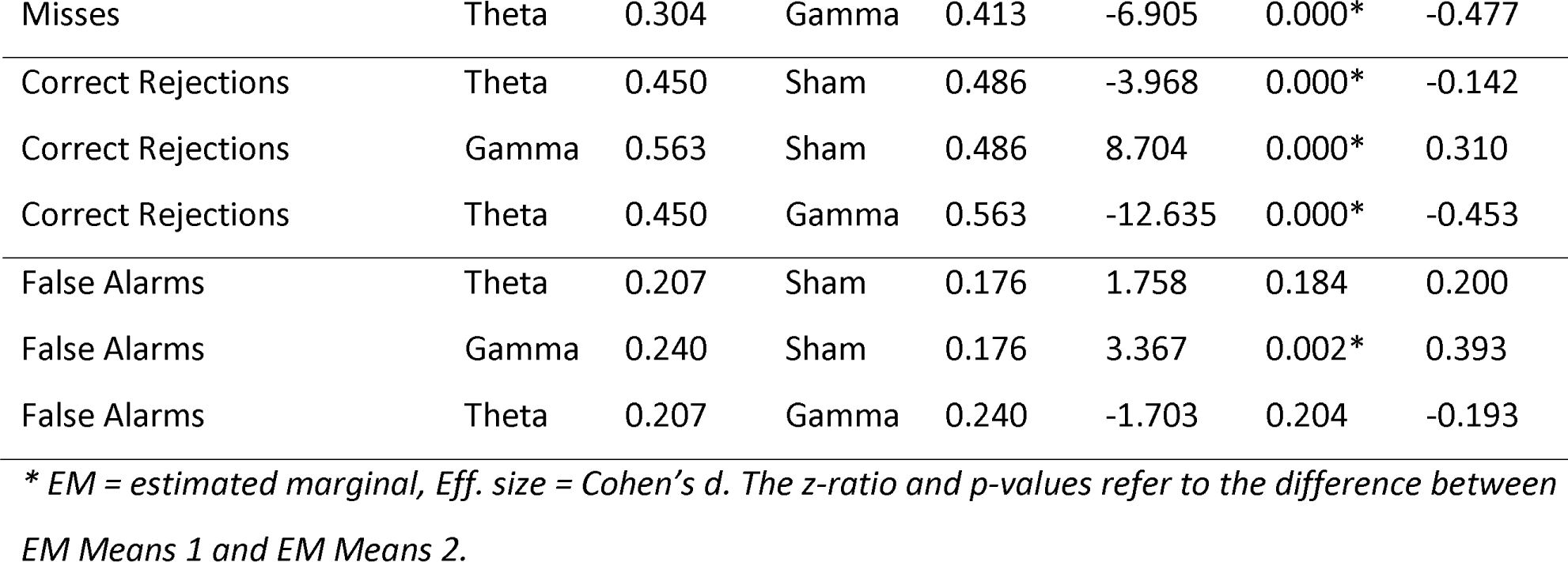
Item Memory Confidence - Post-hoc Tests.

For source memory accuracy and confidence, stimulation had no significant influence on the predicted probability of making a correct (see Table 5) nor a high-confidence decision (see Table 6). However, there was a marginally significant 2-way interaction between ‘Theta tACS’ and ‘Accuracy’ on confidence and a marginally significant 2-way interaction between ‘Gamma tACS’ and ‘Confidence’ on accuracy.

**Table 5.** Source Memory Accuracy – tACS Model.

|  | Estimate | z-value | p-value |
| --- | --- | --- | --- |
| Intercept | 0.791 | 11.681 | 0.000* |
| Gamma tACS | 0.500 | -0.019 | 0.985 |
| Theta tACS | 0.515 | 1.345 | 0.179 |
| Confidence | 0.255 | -31.706 | 0.000* |
| Source | 0.451 | -10.781 | 0.000* |
| Gamma tACS * Confidence | 0.481 | -1.661 | 0.097<sup>+</sup> |
| Theta tACS * Confidence | 0.516 | 1.372 | 0.170 |

**Table 6.**
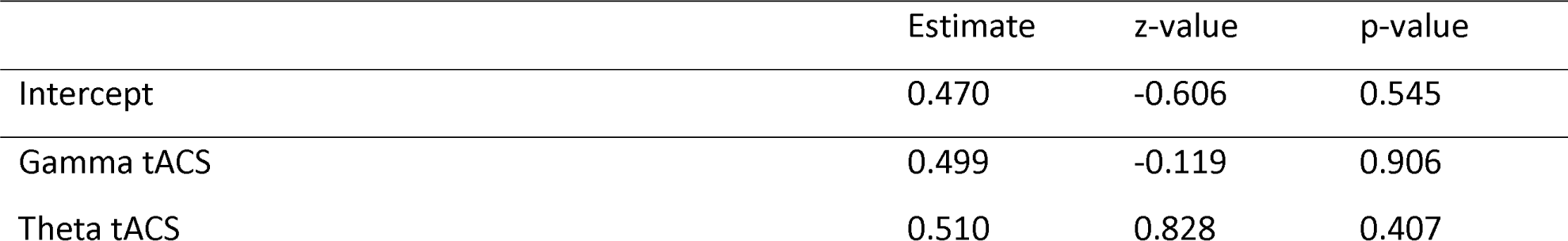
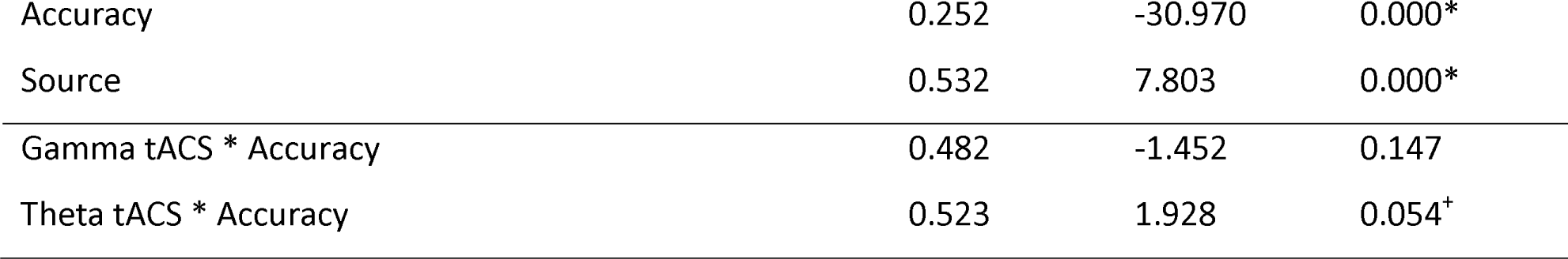
Source Memory Confidence – tACS Model.

|  | Estimate | z-value | p-value |
| --- | --- | --- | --- |
| Intercept | 0.470 | -0.606 | 0.545 |
| Gamma tACS | 0.499 | -0.119 | 0.906 |
| Theta tACS | 0.510 | 0.828 | 0.407 |
| Accuracy | 0.252 | -30.970 | 0.000* |
| Source | 0.532 | 7.803 | 0.000* |
| Gamma tACS * Accuracy | 0.482 | -1.452 | 0.147 |
| Theta tACS * Accuracy | 0.523 | 1.928 | 0.054<sup>+</sup> |

To summarize, during theta tACS participants are more likely to make high-confident hits and less likely to make high-confident correct rejections, while during gamma tACS all responses are more likely to be high-confident responses.^1^ These effects are exclusive to item memory, as we did not find evidence for any significant stimulation effects on source memory.

### 3.3. Predicting individual tACS effects from EEG data

To further explore the tACS effects on item memory, we used the EEG collected during session 1 to explore which EEG correlates can predict the subsequent stimulation effects during sessions 2-4. As tACS effects (see previous section) mainly pertained to high-confident correct responses, and these are also the most relevant, we included only those trials in these models.

The first model looked at theta-related EEG components which can predict the theta tACS effect on high-confident hits. This ‘theta tACS effect’ was defined as the difference score of the proportion of high-confident hits in the theta tACS and sham tACS conditions. This model showed that retrieval-related parietal power, parietal peak deviation, phase synchronization between frontal and parietal channels, and frontal and parietal phase-amplitude coupling were able to predict the stimulation effect on high-confident hits (see Table 7). More specifically, it appears that participants with lower parietal theta power, are predicted to show a larger theta tACS effect on high-confident hits. In addition, a parietal theta peak further away from the stimulation frequency (4 Hz) increases the theta tACS effect. Furthermore, the greater the phase synchronization between frontal and parietal channels, the stronger the theta tACS effect. Lastly, more phase-amplitude coupling between theta and gamma oscillations showed a negative relationship with the theta tACS effect in frontal channels, and a positive relationship with the theta tACS effect in parietal channels.

**Table 7.** High-Confident Hits – Theta EEG/tACS Model.

|  | Estimate | t-value | p-value |
| --- | --- | --- | --- |
| Intercept | -0.018 | -0.365 | 0.718 |
| Encoding |  |  |  |
| Frontal Power | 0.043 | 1.779 | 0.085<sup>+</sup> |
| Parietal Power | -0.012 | -0.523 | 0.605 |
| Frontal Peak Deviation | -0.029 | -1.456 | 0.155 |
| Parietal Peak Deviation | 0.001 | 0.078 | 0.938 |
| Phase Synchronization | -0.027 | -1.517 | 0.139 |
| Frontal Phase-Amplitude Coupling | -0.006 | -0.325 | 0.747 |
| Parietal Phase-Amplitude Coupling | -0.06 | -1.917 | 0.065<sup>+</sup> |
| Retrieval |  |  |  |
| Frontal Power | 0.049 | 1.891 | 0.068 |
| Parietal Power | -0.053 | -2.266 | 0.031<sup>*</sup> |
| Frontal Peak Deviation | -0.039 | -1.662 | 0.107 |
| Parietal Peak Deviation | 0.105 | 4.529 | 0.000<sup>*</sup> |
| Phase Synchronization | 0.075 | 3.614 | 0.001<sup>*</sup> |
| Frontal Phase-Amplitude Coupling | -0.059 | -2.552 | 0.016<sup>*</sup> |
| Parietal Phase-Amplitude Coupling | 0.092 | 2.894 | 0.007<sup>*</sup> |

The second model looked at the theta-related EEG components which can predict theta tACS effects on high-confident correct rejections. This ‘theta tACS effect’ was defined as the difference score of the proportion of high-confident correct rejections in the theta tACS and sham tACS conditions. This model showed that only retrieval-related phase synchronization between frontal and parietal channels has a positive relationship with the subsequent theta tACS effect (see Table 8).

**Table 8.** High-Confident Correct Rejections – Theta EEG/tACS Model.

|  | Estimate | t-value | p-value |
| --- | --- | --- | --- |
| Intercept | -0.006 | -0.149 | 0.883 |
| Retrieval |  |  |  |
| Frontal Power | -0.005 | -0.144 | 0.886 |
| Parietal Power | 0.054 | 1.636 | 0.11 |
| Frontal Peak Deviation | -0.035 | -1.312 | 0.197 |
| Parietal Peak Deviation | 0.033 | 1.403 | 0.168 |
| Phase Synchronization | 0.052 | 2.584 | 0.013* |
| Frontal Phase-Amplitude Coupling | 0.004 | 0.181 | 0.857 |
| Parietal Phase-Amplitude Coupling | -0.011 | -0.474 | 0.638 |

The third model explored the gamma-related EEG components which can predict gamma tACS effects on high-confident hits. This ‘gamma tACS effect’ was defined as the difference score of the proportion of high-confident hits in the gamma tACS and sham tACS conditions. This model did not show any EEG components that had a statistically significant influence on the subsequent stimulation effect (see Table 9). However, the significance of the intercept may be of note here, as this reflects the situation where all the predictors in the model have a value of zero. Here, the predictors ‘power’, ‘phase synchronization’, and ‘phase-amplitude coupling’ are mean-centered, making a value of zero reflecting their mean score. The predictor ‘peak deviation’ is not mean-centered, so a value of zero reflects zero difference between the EEG frequency with the highest power in the gamma range, and the gamma tACS frequency, i.e., an EEG gamma peak frequency that matches the 50 Hz tACS frequency. Therefore, the significant intercept suggests that when averaging out the other predictors, a significant gamma tACS effect is expected when the EEG gamma peak of an individual matches the tACS frequency. However, given that the ‘peak deviation’ predictors are not significant, there is no evidence for a linear relationship between EEG gamma peak deviation and the gamma tACS effect.

**Table 9.** High-Confident Hits – Gamma EEG/tACS Model.

|  | Estimate | t-value | p-value |
| --- | --- | --- | --- |
| Intercept | 0.131 | 3.346 | 0.002* |
| Encoding |  |  |  |
| Frontal Power | -0.019 | -0.847 | 0.403 |
| Parietal Power | 0.029 | 1.423 | 0.164 |
| Frontal Peak Deviation | -0.019 | -1.149 | 0.258 |
| Parietal Peak Deviation | -0.021 | -1.083 | 0.286 |
| Phase Synchronization | -0.024 | -1.373 | 0.178 |
| Frontal Phase-Amplitude Coupling | 0.002 | 0.13 | 0.897 |
| Parietal Phase-Amplitude Coupling | 0.027 | 0.909 | 0.369 |
| Retrieval |  |  |  |
| Frontal Power | -0.021 | -0.821 | 0.417 |
| Parietal Power | 0.017 | 0.602 | 0.551 |
| Frontal Peak Deviation | -0.016 | -0.837 | 0.408 |
| Parietal Peak Deviation | -0.025 | -1.161 | 0.253 |
| Phase Synchronization | 0.018 | 0.932 | 0.358 |
| Frontal Phase-Amplitude Coupling | -0.006 | -0.306 | 0.761 |
| Parietal Phase-Amplitude Coupling | -0.011 | -0.38 | 0.706 |

The fourth model explored the gamma-related EEG components which can predict the gamma tACS effect on high-confident correct rejections. This ‘gamma tACS effect’ was defined as the difference score of the proportion of high-confident correct rejections in the gamma tACS and sham tACS conditions. This model did not show any EEG components that had a statistically significant influence on the subsequent stimulation effect (see Table 10).

**Table 10.**
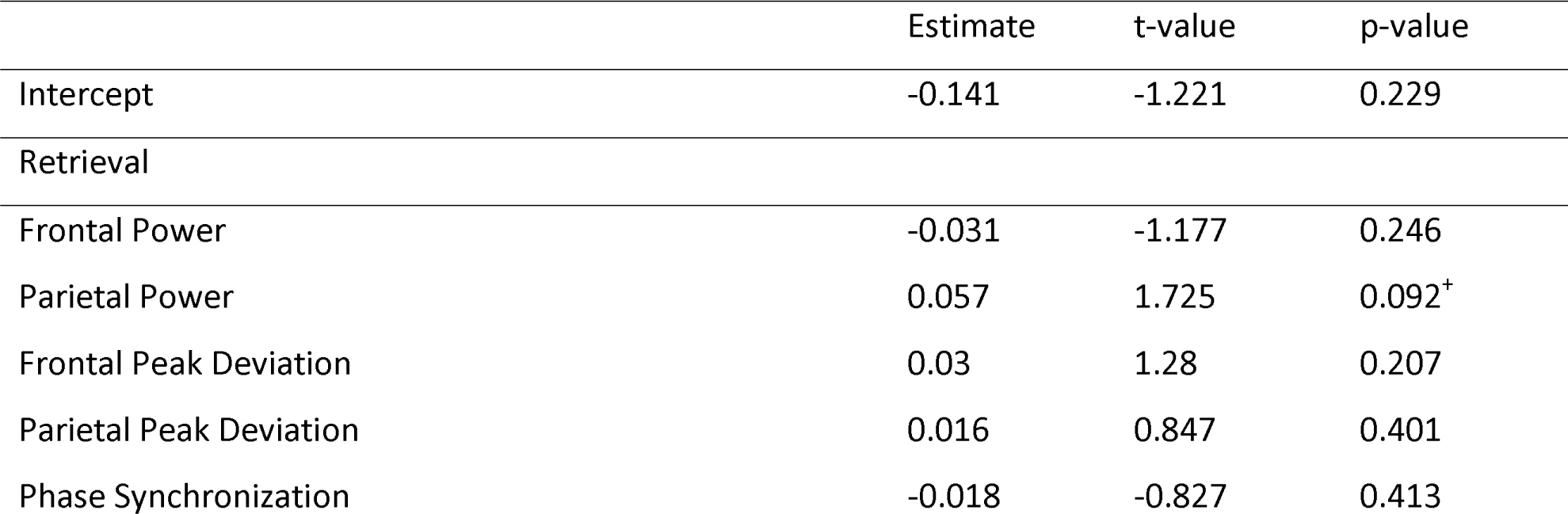
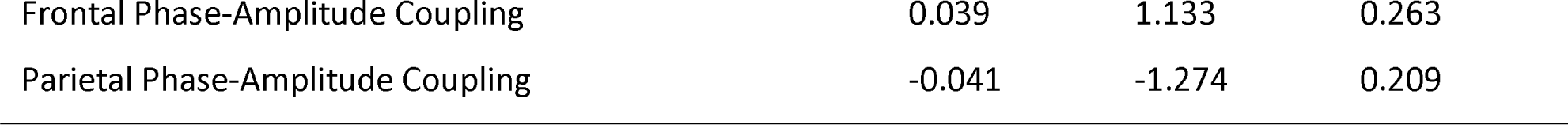
High-Confident Correct Rejections – Gamma EEG/tACS Model.

|  | Estimate | t-value | p-value |
| --- | --- | --- | --- |
| Intercept | -0.141 | -1.221 | 0.229 |
| Retrieval |  |  |  |
| Frontal Power | -0.031 | -1.177 | 0.246 |
| Parietal Power | 0.057 | 1.725 | 0.092<sup>+</sup> |
| Frontal Peak Deviation | 0.03 | 1.28 | 0.207 |
| Parietal Peak Deviation | 0.016 | 0.847 | 0.401 |
| Phase Synchronization | -0.018 | -0.827 | 0.413 |
| Frontal Phase-Amplitude Coupling | 0.039 | 1.133 | 0.263 |
| Parietal Phase-Amplitude Coupling | -0.041 | -1.274 | 0.209 |

To summarize, the results above indicate that only retrieval-related EEG components were able to predict stimulation effects during subsequent tACS. Specifically, oscillatory synchronization measures were able to predict individual differences in subsequent theta tACS effects. In addition, less parietal power, or an oscillatory peak further away from the stimulation frequency seemed to be beneficial for the effect of theta tACS on high-confident hits. For gamma tACS, there were no EEG markers that could predict subsequent gamma tACS effect on behaviour.

## 4. Discussion

This combined EEG and tACS study explored the memory-related behavioural modulation of theta and gamma tACS, and subsequently used EEG markers to predict individual differences in stimulation effects. A source memory task which incorporated confidence ratings was used to investigate the direct involvement of theta and gamma oscillations on item and source memory. A model-based approach was utilized to (1) investigate the effects of frontoparietal theta and gamma tACS on memory accuracy and confidence and (2) explore the EEG components that can predict individual differences in the efficacy of this brain stimulation.

When looking at the effects of theta tACS on memory performance, we see that theta tACS affects both item memory accuracy and item memory confidence in a coherent way. Specifically, theta tACS only significantly increases the probability of a correct response for old items with high-confident responses. Likewise, theta tACS only significantly increased the probability of making a high-confident response for hits. In addition, theta tACS also seems to have a selective negative impact on memory confidence, as it reduced the probability of high-confident responses for correct rejections. Together this suggests that frontoparietal theta tACS can enhance objective and subjective memory in a specific way, by increases the probability of high-confident hits, but it can also reduce subjective novelty detection, by decreasing the probability of high-confident correct rejections. As theta tACS effects were all related to item memory confidence, this stimulation had a greater effect on item than source memory accuracy and confidence. We based our hypothesis regarding source memory mainly on EEG findings showing an increase in theta power during successful source memory (Addante et al., 2011; Gruber et al., 2008; Guderian & Duzel, 2005; Herweg et al., 2016). While two theta otDCS studies found an effect on source memory (Mizrak et al., 2018; Vulic et al., 2021), these results were not replicated in a previous tACS study (Wynn et al., 2020a) and the current study. This indicates that source memory might be more responsive to direct current as compared to alternating current stimulation. Although theta oscillations had already been linked to decision-making confidence (Jacobs, Hwang, Curran, & Kahana, 2006; Selimbeyoglu, Keskin-Ergen, & Demiralp, 2012; Soutschek, Moisa, Ruff, & Tobler, 2021; Wischnewski & Compen, 2022; Wokke, Cleeremans, & Ridderinkhof, 2017), a recent EEG study was one of the first to look at the unique contribution of theta power to source accuracy and source confidence, and found that theta power was significantly linked to confidence, but not accuracy (Wynn et al., 2024). This results in combination with the current findings could indicate that theta oscillations might not play a direct role in source memory accuracy, but that this effect is mediated through confidence. The same may be concluded for item memory, given that the effects seem to be specific to accurate and high-confident responses, indicating an interaction between accuracy and confidence. In item memory, theta oscillations may be specifically involved in the evaluation of retrieved information and the subjective confidence feeling produced by this. When presented with a memory cue, theta tACS might increase the weighting of the relevance and accuracy of retrieved information from memory, leading to more confident hits and more doubt in correct rejections. For hits this would lead to more confidence in the relevant and correct information, and for correct rejections this would lead to less confidence in the novelty of the irrelevant and/or incorrect information retrieved.

We used data from the first EEG-only session to find markers that could predict subsequent tACS effects. When trying to elucidate individual differences in the theta tACS effect on high-confident responses, we looked at EEG components reflecting various aspects of endogenous theta oscillations in high-confident correct response trials. We used EEG data from both encoding and retrieval but found no encoding components that showed a relationship to the subsequent theta tACS effects, only retrieval components. Given that our tACS aimed to strengthen frontoparietal communication, it is noteworthy that the only EEG marker that showed a positive relationship with the theta tACS effect on both high-confident hits and correct rejections was frontoparietal phase synchronization. In other words, participants that show a greater endogenous frontoparietal phase synchronization, are predicted to have a larger faciliatory theta tACS effect on high-confident hits and correct rejections. This indicates that our theta tACS protocol will have a greater effect on individuals who inherently have a higher frontoparietal synchrony. Our other measure of synchrony, phase-amplitude coupling between theta and gamma oscillations, was also a significant predictor of the theta tACS effect, but on high-confident hits only. It has been proposed that NIBS preferentially modulates neuronal networks that are already activated, for example due to the current task-related processes (Bikson, Name, & Rahman, 2013). Therefore, it seems like our tACS protocol, which was aimed to stimulate the frontoparietal network, seems to be specifically effective for people with a higher baseline functional connectivity during memory retrieval. In addition, we found that the theta tACS effect on high-confident hits is predicted to be greater for individuals with lower parietal power and an endogenous parietal theta peak frequency further away from the stimulation frequency (4 Hz). We anticipated that a closer match between the endogenous theta peak frequency and the externally applied tACS frequency would be optimal, but these results show the opposite pattern. This in combination with the negative relationship between parietal theta power and the theta tACS effect might indicate that less endogenous parietal power at the tACS frequency might enable bigger behavioural effects. It has been reported before that low theta (3-4 Hz) oscillations show a greater correlation with memory confidence, as compared to high theta (5-7 Hz) (Wynn et al., 2019; Wynn et al., 2020a). Therefore, if low theta is directly related to the probability of high-confident memory retrieval, it follows that promoting 4 Hz brain oscillations with externally applied tACS would be the most beneficial for participants utilizing this mechanism to a lesser extent intrinsically.

When we shift our attention to the effects of gamma tACS on memory performance, we see that gamma tACS only affected item memory confidence. This gamma tACS effect was found in all memory responses, regardless of memory status or accuracy. This suggests that frontoparietal gamma tACS can influence decision-making processes, in this case memory-related ones. We initially hypothesized similar yet weaker effects of gamma tACS, as compared to theta tACS. However, our results indicate that theta tACS seemed to specifically affect memory confidence in correct responses, whereas gamma tACS affected (memory-related) decision-making. Gamma tACS might specifically influence evidence accumulation during memory retrieval and the subsequent subjective memory experience. Given that gamma oscillations have been linked to evidence accumulation in various task that require a decision (Donner, Siegel, Fries, & Engel, 2009; Polanía, Krajbich, Grueschow, & Ruff, 2014; Solway, Schneider, & Lei, 2022), it is currently unclear whether our current gamma tACS findings are specific to memory-related decision-making, or a general decision-making process. A previous study investigating decision-making metacognition, compared theta and gamma tACS and found that both stimulations increased the link between decision confidence and accuracy (Soutschek et al., 2021). However, this study did not directly assess the effects of gamma tACS on decision-making confidence. Therefore, further research is needed to determine whether the gamma tACS findings we report here are specific to memory-related confidence or to general decision-making confidence. It is of note that gamma tACS also increased memory confidence in incorrect responses (misses and false alarms), suggesting a shift in confidence bias that was not influenced by the accuracy of the decision.

When trying to elucidate individual differences in the gamma tACS effect on high-confident responses, we looked at EEG components reflecting various aspects of endogenous gamma oscillations in high-confident correct response trials. We found no EEG components that showed a relationship to the subsequent gamma tACS effects. This could be due to the lower signal-to-noise ratio in the gamma frequency band when measured with EEG, due to EMG artifacts, making it more difficult to detect small effects (Muthukumaraswamy & Singh, 2013). Regardless of the underlying reason, our results indicate that the EEG components investigated are not able to predict the efficacy of gamma tACS in individuals.

In conclusion, frontoparietal theta tACS can directly influence memory confidence in correct responses, possibly by lowering the “confidence” threshold when evaluating information retrieved from memory. Stimulating the frontoparietal memory network in this way seems to be specifically effective in individuals with greater endogenous frontoparietal connectivity and less endogenous low theta power during memory retrieval. In addition, frontoparietal gamma tACS can directly influence memory confidence, regardless of memory status or accuracy, possibly by shifting the memory confidence bias. EEG could not be used to predict individual differences in this gamma tACS effect.

## Acknowledgements

This work was supported by the National Institutes of Health (NIH) grant R15MH114190. We would like to thank Brandon Lee, Christopher Townsend, and Xiaochen Zheng.

## Footnotes

1 It is important to note that visual inspection of Figure 3 suggested the presence of some data points that could have a disproportional effect on the statistical analysis pertaining to the tACS effect on memory performance. We opted to include all participants in this analysis as their overall task performance did not warrant any exclusion and our task design is only fully balanced when all participants are included. We therefore have no reason to believe these data points are not part of the natural variability of the data. When excluding the four participants that were over three standard deviations away from the mean in the stimulation effects on high-confident hits and correct rejections (shown in Figure 3) the overall results were comparable. The only result that changed was the theta tACS effect on correct rejections; the direction of this effect was still negative, yet not significant anymore. This indicates that the observed effects are robust to the inclusion or exclusion of these outliers, except for the specific theta tACS effect on correct rejections.

